# Subpopulations of cancer-associated fibroblasts expressing fibroblast activation protein and podoplanin in non-small cell lung cancer are a predictor of poor clinical outcome

**DOI:** 10.1101/2022.09.28.509919

**Authors:** Layla Mathieson, Lilian Koppensteiner, Samuel Pattle, David A Dorward, Richard O’Connor, Ahsan Akram

## Abstract

Cancer-associated fibroblasts (CAFs) are the dominant cell type in the stroma of solid organ cancers, including non-small cell lung cancer (NSCLC). Fibroblast heterogeneity is widely recognised in many cancers, with subpopulations of CAFs being identified and potentially being indicative of prognosis and treatment efficacy. Here, the subtypes displayed by CAFs isolated from human NSCLC resections are initially identified by flow cytometry, using the markers FAP, CD29, αSMA, PDPN, CD90, FSP-1 and PDGFRβ, showing five distinct subpopulations, CAF-S1-S5. Our findings show that when comparing fibroblasts from tumour tissue with that from adjacent lung tissue, CAF-S2 and CAF-S3 are found in the normal tissue and marker expression suggests a less activated phenotype whereas CAF-S1, CAF-S4 and CAF-S5 are predominantly found in the tumour tissue and are positive for a combination of markers of fibroblast activation. We focus on these subtypes most associated with fibroblast activation, primarily focussing on a previously unreported CAF-S5 subtype, and comparing to the previously identified CAF-S1. Both these subsets express FAP and PDPN as markers of fibroblast activation, but CAF-S5 lacks expression of the common activation marker αSMA. The spatial relevance of these subtypes in a cohort of 163 NSCLC patients was then investigated by multiplex immunofluorescence on a tumour micro-array of patient samples, revealing CAF-S5 are found further from tumour regions than CAF-S1. To understand the functional role of CAF-S5, scRNA sequencing data was used to compare the subset to the previously identified CAF-S1, finding that CAF-S5 displays an inflammatory phenotype, whereas CAF-S1 displays a contractile phenotype. We demonstrate that presence of either the CAF-S1 or CAF-S5 subtype is correlated to worse survival outcome in NSCLC, highlighting the importance of the identification of CAF subtypes in NSCLC.

## Introduction

Lung cancer is the leading cause of cancer death globally (1) and non-small cell lung cancer (NSCLC) accounts for approximately 85% of cases (2). Current NSCLC therapies are often unsuccessful, with drug resistance leading to treatment failure and disease progression (3). The tumour stroma plays a role in this resistance to therapy and has emerged as an important target for therapies to combat cancers such as NSCLC (4–7). The most common cell type of the tumour stroma is the cancer-associated fibroblast (CAF) (8). In healthy tissue, fibroblasts are a quiescent structural component of the ECM, and become activated in response to wound signals. In their activated state they produce ECM components and engage in crosstalk with immune cells to promote wound healing. After the healing process is complete, fibroblasts return to a quiescent state and excess fibroblasts are removed by apoptosis (9). CAFs on the other hand, are found in an irreversibly activated state. They have been found to have an enhanced migratory phenotype over normal activated fibroblasts, a greater proliferative ability and an enhanced secretome (10). CAFs have been found to play a role in immune evasion, metastasis, invasion, angiogenesis and resistance to drug treatment (6,11,12).

Several studies have shown that CAFs represent a heterogeneous population composed of functionally distinct subtypes (6,13–18). The phenotype of these subtypes has been characterised in some solid organ malignancies, including breast, ovarian, pancreatic and lung cancers (14,17–21). Markers frequently used to distinguish these subtypes include α-smooth muscle actin (αSMA), fibroblast activation protein (FAP), podoplanin (PDPN), integrin β1 (CD29) and fibroblast-specific protein-1 (FSP-1). Two key subtypes of note, previously termed CAF-S1 and CAF-S4 in the literature, have been identified in several studies. CAF-S1 display a FAP^hi^ phenotype associated with adhesion, wound healing and immunosuppression while CAF-S4 which are FAP^low/negative^, express higher levels of αSMA and are associated with invasion and metastasis(7,14,17,22–25). Spatially, CAF-S1 have been found in closer proximity to cancer cells. The presence of these subtypes can also indicate prognosis, with CAF-S1 and CAF-S4 being found to promote metastases, and CAF-S1 being an indicator or distant relapse in luminal breast cancer (17).

Here, we investigate CAF subtypes present in NSCLC, identifying five subtypes using commonly used CAF markers. We focus on the previously unreported CAF-S5 subtype, identified primarily by the expression of FAP and PDPN but lacks expression of αSMA. We compare the spatial location of CAF-S5 to the previously defined CAF-S1 subtype, and investigate the correlation of these subtypes to survival outcome.

## Methods

### Ethics Statement

Cancer tissue was obtained following approval by NHS Lothian REC and facilitated by NHS Lothian SAHSC Bioresource (REC No: 15/ES/0094). All participants provided written informed consent. NSCLC tissues lung samples (cancer and non-cancerous lung) were collected from patients undergoing surgical resection with curative intent. The tissue microarray was approved NHS Lothian REC and facilitated by NHS Lothian SAHSC Bioresource (REC No: 15/ES/0094) and approved by delegated authority granted to R&D by the NHS Lothian Caldicott Guardian (Application number CRD19031)

### NSCLC Patient Sample Processing

CAFs were isolated from NSCLC patient samples as previously described (26). Briefly, tissue samples were minced with forceps and incubated for an hour in prewarmed RPMI media (Gibco) containing collagenase IV [2 mg/ml] (Sigma) and DNase [0.2 mg/ml] (Sigma). Samples were centrifuged at 300 g for 5 minutes and red blood cells were lysed from samples using RBC lysis buffer (Roche) in 5 ml for 10 minutes at room temperature. Cells were washed in plain RPMI media and then counted in preparation for staining.

### Flow Cytometry Sample Preparation

Cells were collected in suspension post digest and at each passage and were stained with a live/dead marker Zombie UV (1:1000, Biolegend) for 30 min at room temperature in DPBS (Gibco). Cells were then washed and incubated with FC blocker for 10mins and then stained with surface marker antibodies (EpCAM, CD45, CD31, FAP, CD29, Podoplanin and PDGFRβ, see Table 1 for details) for 20 mins at 4°C in DPBS supplemented with 2% FBS. After washing cells were fixed with Cytofix fixation buffer for 20 mins at 4°C. Cells were then washed in Perm/Wash buffer and centrifuged at 300g for 5 mins. Intracellular antibodies (αSMA and FSP-1) were diluted in Perm/Wash buffer then added to cells and incubated in the dark for 30 mins at 4°C. After washing, cells were stored in DPBS with 2% FBS overnight at 4°C before data acquisition on a LSR6Fortessa analyser (BD Biosciences). Compensation was carried out using single stain control UltraComp eBeads (Invitrogen) and isotype control samples were stained using the control antibodies shown in Table 1.

**Table 1:** Antibodies used for flow cytometry staining.

| Marker | Colour | Supplier | ul/test | Catalogue No. | Isotype | ul/test | Iso Catalogue No. |
| --- | --- | --- | --- | --- | --- | --- | --- |
| CD45 | BV605 | BioLegend | 5 | 368524 | IgG1 M | 5 | 400161 |
| CD31 | BV605 | BioLegend | 5 | 303122 | IgG1 M | 5 | 400161 |
| EpCAM | BV605 | BioLegend | 5 | 324224 | IgG2a M | 5 | 400349 |
| CD90 | VioBlue | Miltenyi | 2 | 130-119-890 | IgG1 M | 2 | 130-113-767 |
| Zombie | UV | BioLegend | 1 | 423108 | NA | NA | NA |
| FAP | APC | R&D | 5 | FAB3715A | IgG1 M | 5 | IC002A |
| PDGFRβ | AF594 | R&D | 5 | FAB1263T | IgG1 M | 5 | IC002T |
| CD29 | AF488 | BioLegend | 5 | 303016 | IgG1 M | 5 | 400129 |
| PDPN | APC-Cy7 | BioLegend | 5 | 337030 | IgG2a R | 2.5 | 400524 |
| αSMA | AF750 | R&D | 5 | IC1420S | IgG2a M | 5 | IC 003S |
| FSP-1 | PE | BioLegend | 5 | 370004 | IgG1 M | 5 | 400139 |

### Flow Cytometry Data Analysis

Flow cytometry data was analysed using FlowJo version 10.7.1. Cells were gated to fibroblast populations defined as CD45^-^, EpCAM^-^ and CD31^-^ cells (full gating strategy shown in Fig S1). To reduce file sizes for analysis, fibroblast populations were downsampled to 300 events using the Downsample plugin. Samples containing less than 300 fibroblasts were excluded from analysis. All sample files were then concatenated and from this file UMaps could be generated from the data (27). FlowSOM analysis could then be carried out to determine clusters and was run without defining the number of clusters expected to be unbiased (28). MFIs calculated were the geometric fluorescence intensity.

### Multiplex Immunofluorescence Staining

A TMA was constructed from consecutive patients undergoing surgery with curative intent at a regional thoracic surgery centre over a 2-year period. Following annotation by an experienced thoracic pathologist 1mm cores were taken from tumours and non-cancerous lung for each patient. TMA construct was linked to demographic clinical data and follow up data including both relapse and survival. All patients were treatment naive. TMA slides were deparaffinised in Xylene and rehydrated in a series of ethanol dilutions. Using a Leica Bond automated staining robot; after heat-induced antigen retrieval (HIER) of 30min at 100°C, tissue slides were exposed to multiple staining cycles each including a 30 minute incubation with a protein block (Akoya), 1 hour incubation with the respective primary antibody, 30 minute incubation with the secondary antibody (Akoya), 10 minute incubation with the respective OPAL (Akoya) followed by 20 minute incubation with AR6 buffer (Akoya) at 85°C prior to the next staining cycles and finally stained with fluorescent DAPI (Akoya) for 10 minutes. In between each step, slides were washed with bond wash for 5 minutes.

Primary antibody concentrations and OPAL pairings are shown in Table 2. Antibody-OPAL pairings were assigned based on expected biomarker abundance and expected co-expression.

**Table 2:** Multiplex immunofluorescence antibodies used and their OPAL pairings.

| Primary Antibody | Catalogue No. | Antibody Dilution | OPAL Pairing | OPAL Dilution | Staining Position |
| --- | --- | --- | --- | --- | --- |
| FAP | Ab207178 | 1:100 | OPAL 520 | 1:150 | 1 |
| CD90 | Ab92574 | 1:50 | OPAL 620 | 1:100 | 2 |
| FSP1 | Ab197896 | 1:4000 | OPAL 570 | 1:150 | 3 |
| PDPN | Ab236529 | 1:4000 | OPAL 650 | 1:150 | 4 |
| αSMA | Ab124964 | 1:1000 | OPAL 690 | 1:150 | 5 |
| PanCK | Ab27988 | 1:200 | OPAL 540 | 1:150 | 6 |

### Multiplex Immunofluorescence Imaging

Slides were imaged using a Vectra Polaris. The appropriate exposure time for image acquisition was set for each fluorophore by auto exposing on multiple (5-10) tissue areas per batch. Following fluorescence whole slide scans, regions of interest were selected for multispectral imaging (MSI) at 20x magnification.

### Multiplex Immunofluorescence Image Analysis

MSI images were unmixed in InForm software using representative snapshots of spectral library slides imaged at the same magnification. This also allowed for the isolation of auto fluorescence. Unmixed images were exported and analysed in Qupath (29). Cell detection was performed using StarDist based on a watershed deep-learning algorithm and fluorescent threshold of DAPI nuclear staining (30). Following this, phenotyping was performed in a non-hierarchical manner by creating a composite classifier of single channel classifiers for each stain based on a fluorescent threshold. Ultimately, a machine learning algorithm was trained on multiple images to detect tumour and stroma areas. For each image the counts of the number of cells classified by each combination of markers was calculated and exported for analysis using R.

### Single Cell RNA Sequencing Analysis

Open source data from Lembrechts et al. (20) was analysed using R. The fibroblast data set was downloaded and filtered for fibroblasts that could be defined as CAF-S1 or CAF-S5 using the definitions of the subtypes established by flow cytometry. Fibroblasts were filtered by including those with expression of CD29, PDGFRβ, PDPN and FAP and excluding any that expressed FSP1. The remaining fibroblasts were then determined to be CAF-S1 if they expressed αSMA above 10 counts, and CAF-S5 if they did not express αSMA. A PCA plot of the resulting subset of fibroblasts was created using Orange Data Mining (31). Differential expression analysis was then performed in R using the DESeq2 package (32). The top 100 genes were plotted in a heatmap to assess key differences between the two subtypes and a volcano plot generated using the enhanced volcano package (33).

### Analysis of Survival Data

Survival data was collected for the 163 NSCLC patients whose samples were included in the TMA analysed by multiplex immunofluorescence, where survival was defined as the number of days from surgery to death or follow up. Kaplan-Meier curves were plotted for patients who had fibroblasts of phenotype CAF-S1 or CAF-S5 present (determined in QuPath, described above) above and below the median number of CAFs present in that subtype. Log-rank tests were used to determine significance. Plots were also generated for the markers FAP, PDPN and αSMA, showing survival when these markers are present above or below median expression levels. Analysis was carried out using the survival and survminer packages in R.

### Analysis of TCGA Data

Data for liver hepatocellular carcinoma, pancreatic adenocarcinoma, breast invasive carcinoma and kidney renal clear cell carcinoma was downloaded from https://tcga-data.nci.nih.gov. The surv_cutpoint function in R was used to determine the most significant cut off for expression level correlated to survival for each cancer for the markers FAP, PDPN and αSMA. Using these cut-offs generated patients could be defined as low or high for each marker. Patients were considered to have an overall CAF-S5 like phenotype if they were FAP and PDPN high and αSMA low. The survival of these patients was then compared all other patients by plotting Kaplan-Meier curves as previously described.

## Results

To understand the heterogeneity of CAFs in human NSCLC we first looked at the expression levels of seven CAF markers using flow cytometry (Fig 1A(i)). As no single fibroblast marker exists, fibroblasts were identified as being negative for EpCAM, CD45 and CD31 to exclude epithelial, hematopoietic and endothelial cells respectively (Fig 1A(ii)). Fibroblast markers FAP, CD29, αSMA, PDPN, CD90, FSP1 and PDGFRβ expression levels were determined and compared for tumour and non-cancerous adjacent lung tissue from NSCLC patients (Fig 1B). The markers FAP, CD29, αSMA, PDPN, CD90 and PDGFRβ typically showed elevated expression in tumour compared to non-cancerous lung tissue, whereas FSP1 showed downregulation in tumour compared to non-cancerous lung. Across all markers it was clear that there was significant variance between patients, confirming CAF heterogeneity in NSCLC.

**Figure 1:**
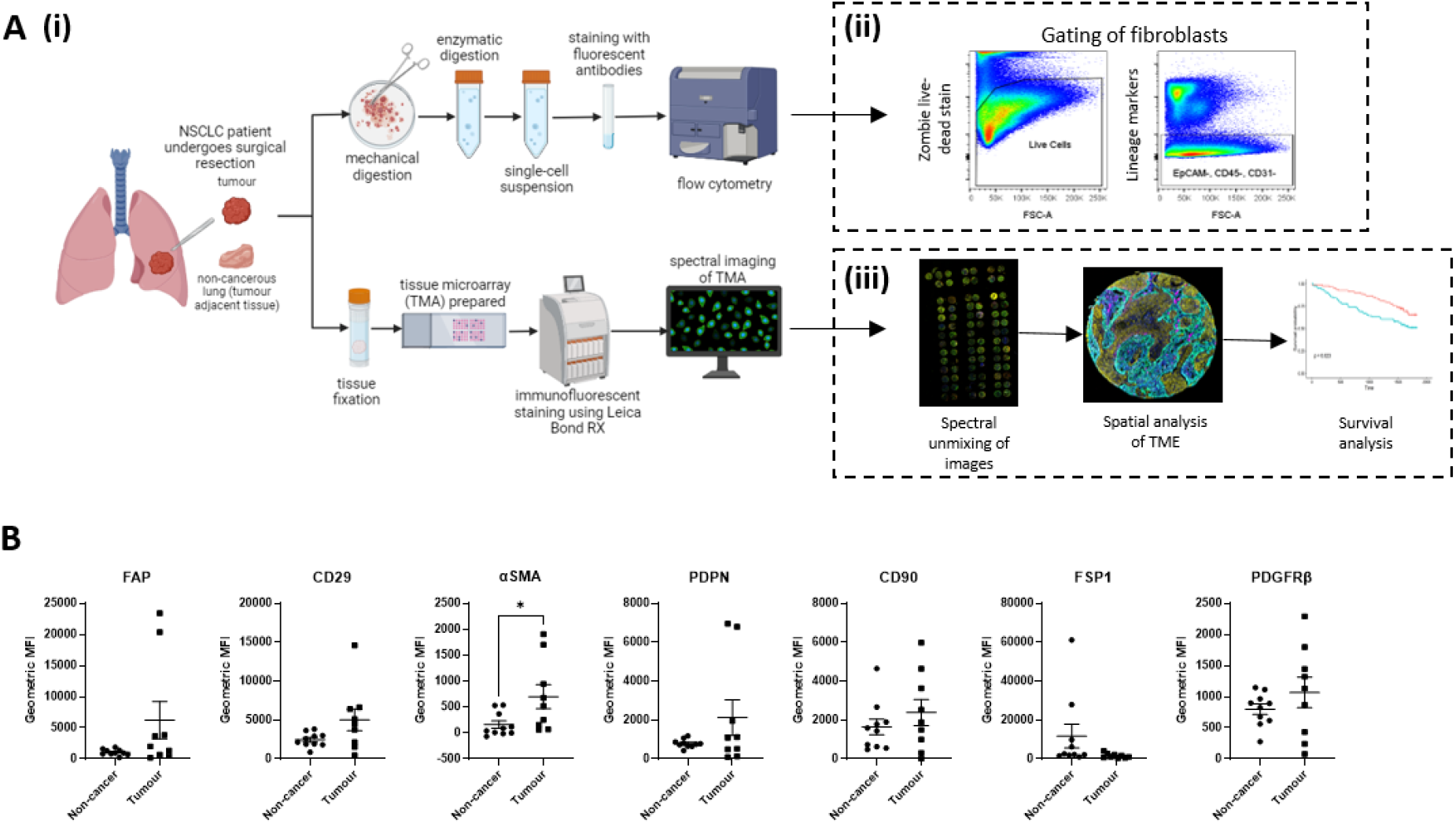
Identifying CAFs in NSCLC by expression of fibroblast markers. (A) The preparation of NSCLC samples for analysis of CAFs from NSCLC patient resections for analysis by flow cytometry (ii) or multiplex immunofluorescence (iii); (B) Expression levels of FAP, CD29, αSMA, PDPN, CD90, FSP1 and PDGFRβ determined by FACS in non-cancerous lung tissue compared to tumour tissue. Individual data points shown (tumour n=9, non-cancer n=10) as well as mean ±SEM. Unpaired t-test, *p<0.05. Images created with Biorender.com.

To further investigate CAF heterogeneity among NSCLC patients, FlowSOM (28) was used to determine phenotypic clusters of CAFs in an unbiased manner. This identified five subsets of CAFs across the samples (Fig 2A), which we named CAF-S1 (pink), CAF-S2 (red), CAF-S3 (green), CAF-S4 (blue) and CAF-S5 (orange) following previous work by other researchers in breast and ovarian cancers (17,18,34). These subsets were best identified according to their expression levels of FAP and αSMA, as the five subsets could be distinctly identified (Fig 2B), whereas when comparing other fibroblast markers it was less clear (Fig 2D). Across nine patient samples, significant heterogeneity of CAFs was found, these subsets were not found to represent a majority of an individual patient, but rather patients exhibited heterogeneity within their CAF population (Fig 2C).

**Figure 2:**
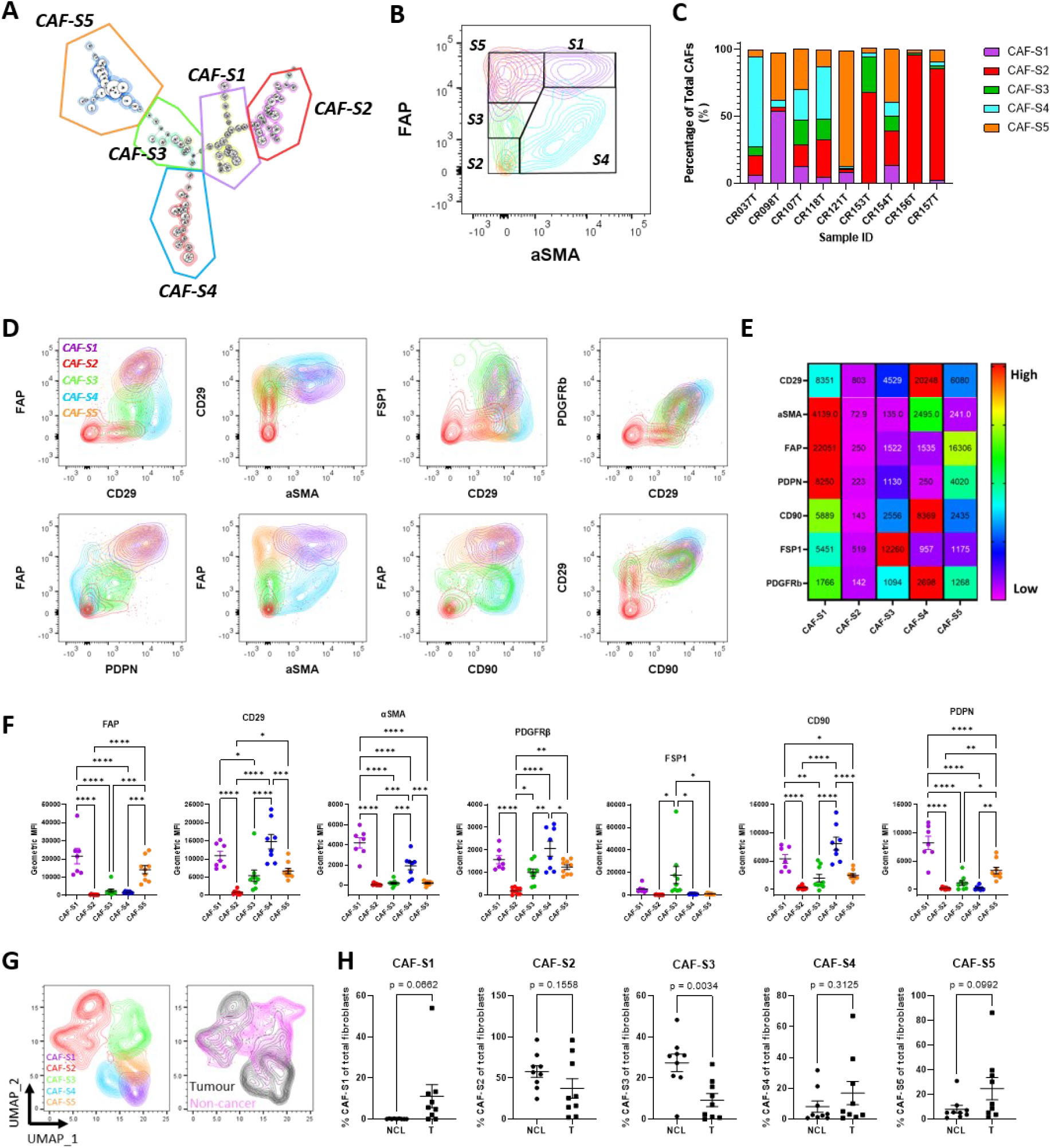
CAF subsets identified in NSCLC. (A) FlowSOM plot showing identification of five CAF subsets in NSCLC, CAF-S1 – S5; (B) Contour plot showing how FAP and αSMA can be used to distinguish CAF subsets in NSCLC; (C) Breakdown of CAF subsets in individual NSCLC samples; (D) Expression profiles of the identified CAF subsets using the different CAF markers; (E) Heat map showing the relative levels of expression of each CAF marker between identified subsets; (F) The expression levels of each marker within each subset. Each point represents geometric MFI of that marker for each sample that contained CAFs of that subset. Stats show Tukey’s multiple comparisons test results, (*p≤0.05, **p ≤ 0.01, ***p ≤ 0.001, ****p ≤ 0.0001); (G) UMAPs showing the clustering of the CAF subsets and the comparison of tumour and non-cancerous fibroblasts showing overlap within some subsets; (H) Comparison of the percentage of each CAF subset present in non-cancerous lung tissue (NCL) with tumour tissue. P-values calculated using unpaired t test.

Comparing the expression levels of each CAF marker within the identified subsets, we classified each subsets expression profile (using Fig 2E&F) as:

CAF-S1: FAP^High^ CD29^Med-High^ αSMA^High^ PDPN^High^ CD90^Med^ FSP1^Low^ PDGFRβ^Med^,

CAF-S2: FAP^Neg^ CD29^Neg-Low^ αSMA^Neg^ PDPN^Neg^ CD90^Neg^ FSP1^Neg^ PDGFRβ^Neg^,

CAF-S3: FAP^Low^ CD29^Med^ αSMA^Neg-Low^ PDPN^Low^ CD90^Low^ FSP1^High^ PDGFRβ^Low^,

CAF-S4: FAP^Neg-Low^ CD29^High^ αSMA^Med^ PDPN^Neg^ CD90^High^ FSP1^Neg^ PDGFRβ^Med-High^ and

CAF-S5: FAP^Med^ CD29^Med^ αSMA^Neg-Low^ PDPN^Med^ CD90^Low^ FSP1^Low^ PDGFRβ^Med^.

Following dimensionality reduction of the data by uniform manifold approximation and projection (UMAP), the fibroblast populations between tumour and non-cancerous samples were compared and it was observed that there was overlap between CAFs and NCL fibroblasts (Fig 2G). Upon investigating the percentage each CAF subset represented of the total fibroblast population in each sample, the proportions across tumour and NCL could be assessed (Fig 2H). This revealed that subsets CAF-S2 and CAF-S3 were more representative of a normal lung fibroblast than a CAF, and hence we focussed on CAF-S1, CAF-S4 and CAF-S5 for analysis in NSCLC tumour samples.

Next, we investigated the spatial location and distribution of CAF subsets in NSCLC by multiplex immunofluorescent (MIF) staining of a tissue microarray (TMA) of 163 tumours. Tumour cores were stained with PanCK to identify tumour regions and the fibroblast makers FAP, PDPN, αSMA, FSP1 and CD90 were used to identify the key CAF subsets identified above as being predominant in tumour tissue: CAF-S1, CAF-S4 and CAF-S5. Using the definitions established by flow cytometry to characterise a profile for each subset as having markers on or off we initially defined subsets as: CAF-S1: FAP^ON^ αSMA^ON^ FSP1^OFF^ CD90^ON^ PDPN^ON^, CAF-S4: FAP^OFF^ αSMA^ON^ FSP1^OFF^ CD90^ON^ PDPN^OFF^, CAF-S5: FAP^ON^ αSMA^OFF^ FSP1^OFF^ CD90^OFF^ PDPN^ON^. This binary classification allowed for classification of individual cells as each subtype.

The MIF results showed clear staining of the fibroblasts markers in only the stromal regions, with the tumour regions stained by PanCK (Fig 3A). As an initial investigation into the staining profile of each fibroblast marker used, the percentage of stromal cells positive for each marker was investigated across disease subtypes. This revealed the level of heterogeneity between patients across subtypes, with the greatest range in expression levels shown in FAP and PDPN expression (Fig 3B). PDPN expression also showed significant difference in expression levels between adenocarcinoma and squamous cell carcinoma, showing higher percentage positivity of PDPN in squamous cell carcinoma patients. It was also observed that staining for CD90 was low, with very few cells classed as CD90^+^ across different classes of NSCLC (Fig 3B). Therefore CD90 was not be used as a marker to characterise the CAF subsets, as differentiation of CAF-S1, CAF-S4 and CAF-S5 remined possible. The final definitions used for the MIF analysis were therefore: CAF-S1: FAP^ON^ αSMA^ON^ FSP1^OFF^ PDPN^ON^, CAF-S4: FAP^OFF^ αSMA^ON^ FSP1^OFF^ PDPN^OFF^, CAF-S5: FAP^ON^ αSMA^OFF^ FSP1^OFF^ PDPN^ON^.

**Figure 3:**
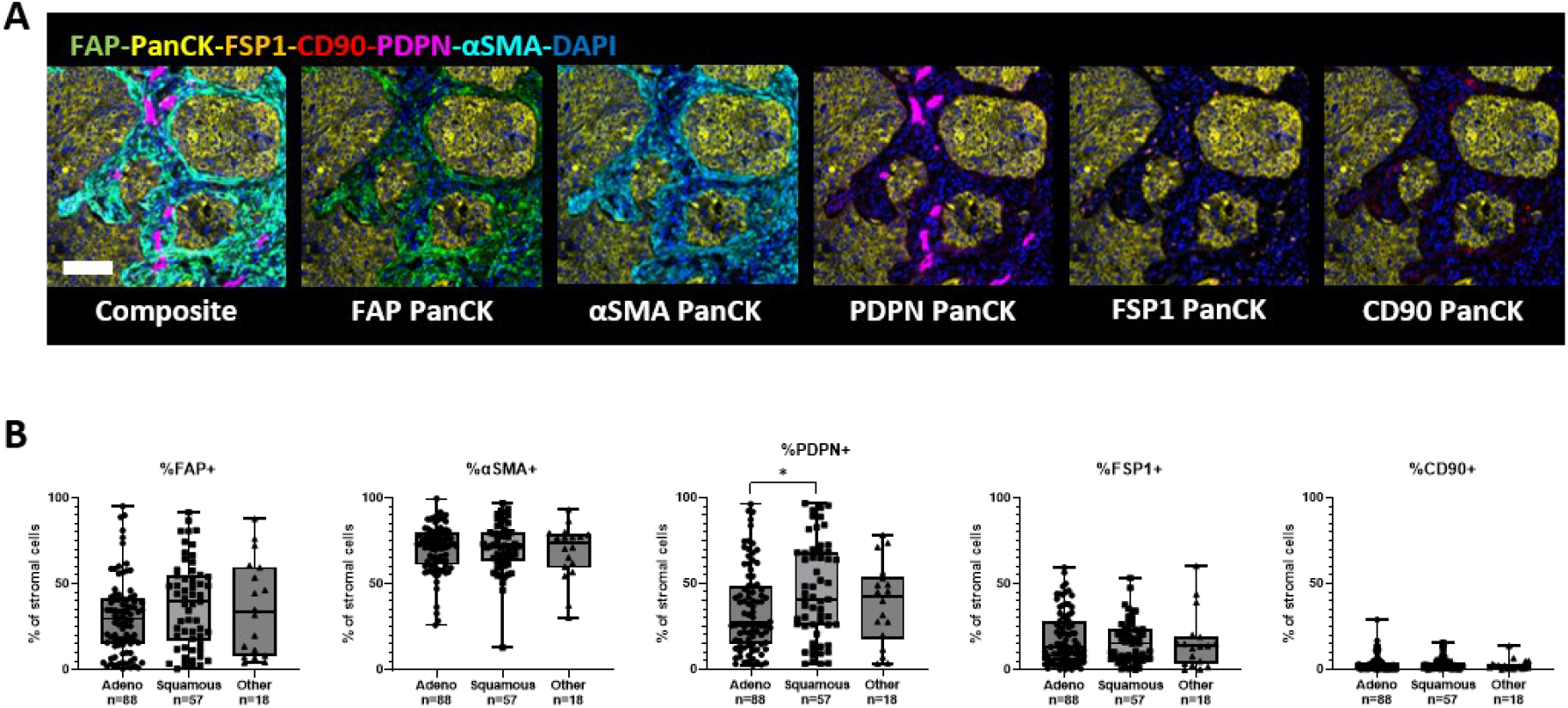
Multiplex immunofluorescence staining of CAFs in NSCLC. (A) Representative images showing the expression pattern of CAF markers FAP, αSMA, PDPN, FSP1 and CD90 relative to cancer cells identified by PanCK staining in a NSCLC tumour sample, scale bar 100um; (B) The percentage of stromal cells positive for CAF markers in different categories of NSCLC. Stats show Tukey’s multiple comparisons test results, *p ≤ 0.05. N=163.

Following segmentation of cells and tissue types in QuPath (Fig 4A), CAFs could be categorised into subsets depending on the markers they expressed. To understand the distribution of the CAF subsets, we investigated whether different subsets dominated in different types of NSCLC by calculating the percentage of stromal cells that were each CAF subset for adenocarcinoma, squamous cell carcinoma and other NSCLC subtypes (Fig 4B). This revealed that CAF-S1 and CAF-S5 were both upregulated in squamous cell carcinoma compared to adenocarcinoma, whereas CAF-S4 was upregulated in adenocarcinoma. This raised questions about the similarities of CAF-S1 and CAF-S5, as they showed the same trend. We first considered whether there was a spatial difference between the two subtypes, as we had previously observed that αSMA staining was dominant near tumour regions (Fig 3A), and the key difference between the two subtypes is the lack of αSMA expression on CAF-S5 compared to CAF-S1. The spatial distribution was quantified by calculating the distance from each CAF to the nearest tumour region (Fig 4C). This showed that CAF-S5 were more likely to be found further from tumour regions than CAF-S1.

**Figure 4:**
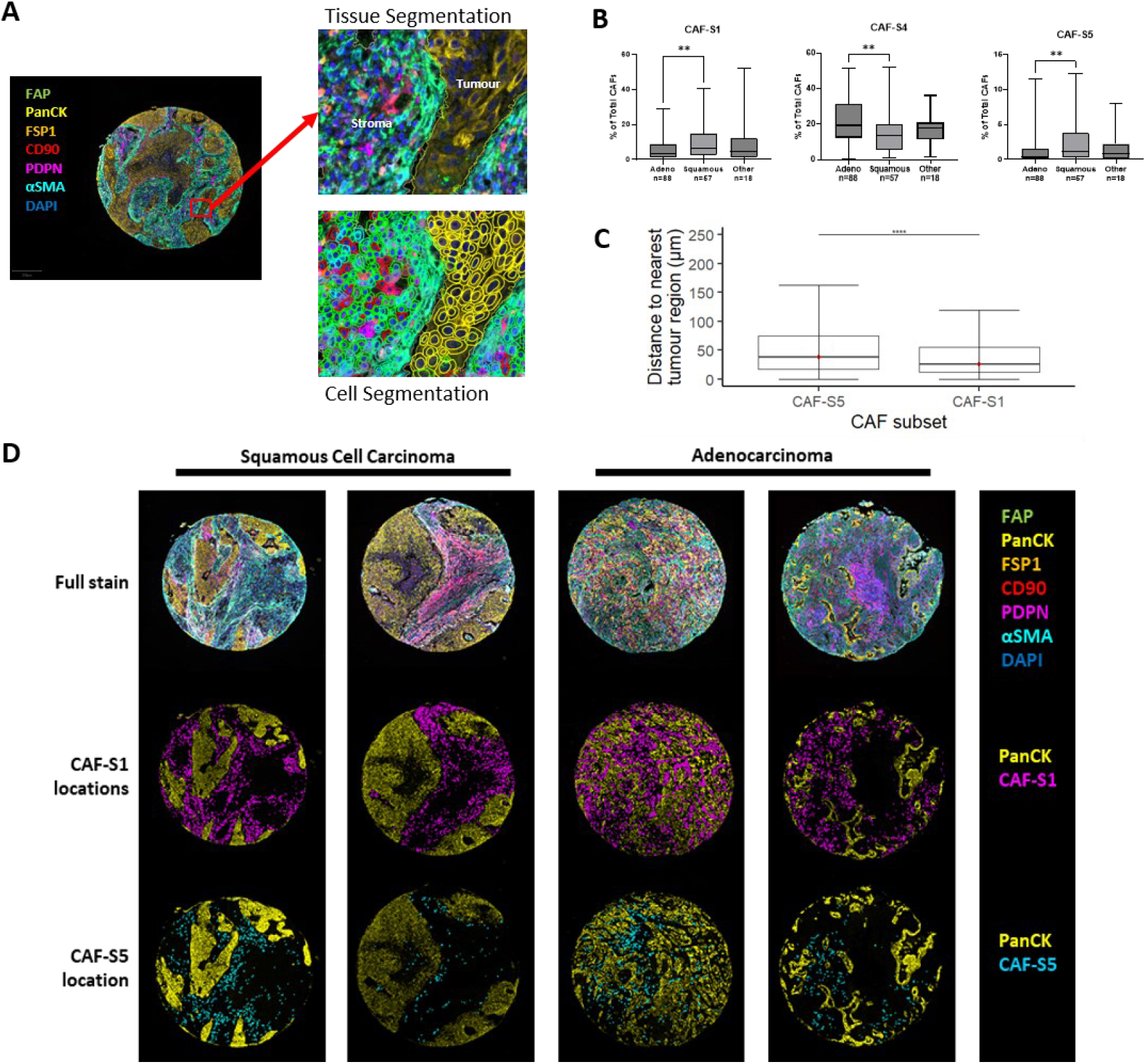
Spatial location of CAF subsets in NSCLC. (A) Segmentation strategy implemented in QuPath to define tissue class as tumour or stroma and to individually segment cells for classification; (B) Quantification of distance of fibroblasts of each class from their nearest tumour region. Data points represent individual fibroblasts from 163 tumour samples; (C) The percentage of stromal cells that are in each CAF subset. Statistics show Tukey’s multiple comparisons test results (*p≤0.05, **p ≤ 0.01); (D) Representative images showing the spatial location of CAF-S1 and CAF-S5 in squamous cell and adenocarcinoma.

To further understand the distinction between CAF-S1 and CAF-S5, single cell RNA sequencing data, available from Lambrechts et al. (20) was analysed to reveal functional differences. Initially, principal component analysis was performed on a selection of CAFs identified using the previously stated definitions to check if clustering of the two subtypes was observed (Fig 5A), which was found to be the case. From our previous results, we know that the main classification difference between CAF-S1 and CAF-S5 is the expression of αSMA, with CAF-S1 highly expressing this and CAF-S5 having negative to low expression levels of it. When plotting a heatmap of the top 100 differentially expressed genes, we observed that CAF-S1 and CAF-S5 do cluster separately, further reassuring that they are distinct subtypes (Fig 5B). When investigating the most downregulated genes in CAF-S5 when compared to CAF-S1 (Fig 5C) we see that the other genes of significance are TAGLN (transgelin), TPM2 (tropomyosin 2), SPARC (secreted protein acidic and cysteine rich) and MYL9 (myosin light chain 9). The upregulated genes are C3 (complement C3), SEPP1 (selenoprotein P), C7 (complement C7) and CLU (clusterin). The results of these analyses suggest that CAF-S1 and CAF-S5 are distinct CAF subtypes.

**Figure 5:**
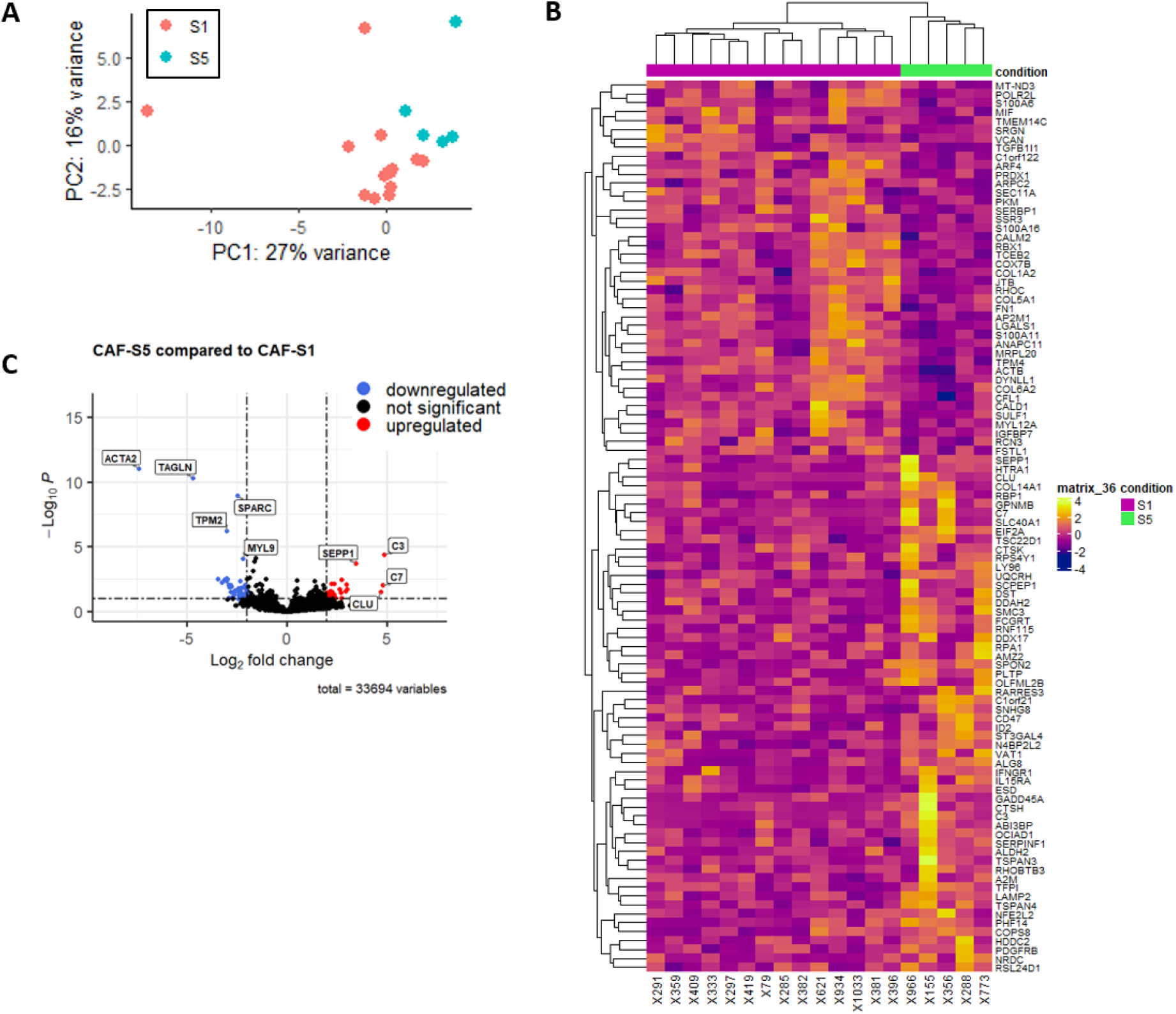
Functional analysis of CAF-S1 and CAF-S5 in NSCLC. (A) PCA plot comparing CAF-S1 and CAF-S5 fibroblasts from Lambrechts et al. RNA Seq data (20) (n = 12 CAF-S1, n = 5 CAF-S5); (B) Volcano plot showing the most significantly up and downregulated genes when comparing CAF-S5 to CAF-S1; (C) Gene sets activated and suppressed in CAF-S5 compare to CAF-S1 following gene set enrichment analysis; (D) KEGG pathways activated and suppressed in CAF-S5 compared to CAF-S1 following KEGG pathway analysis.

Next, we performed survival analysis on our results from 163 NSCLC tumours, looking at if the presence of CAF-S1 and CAF-S5 correlated with recurrence free-survival (RFS) (Fig 6A). This revealed that the presence of CAF-S1 or CAF-S5 was correlated with poor 5-year RFS. As it was evident some patients expressed both CAF-S1 and CAF-S5, survival analysis was performed to compare those that expressed only one of the two subsets above median level with those that expressed both (Fig 6B). This revealed no significant difference between RFS rates of the three groups, with all three demonstrating around 50% RFS probability after 5 years.

**Figure 6:**
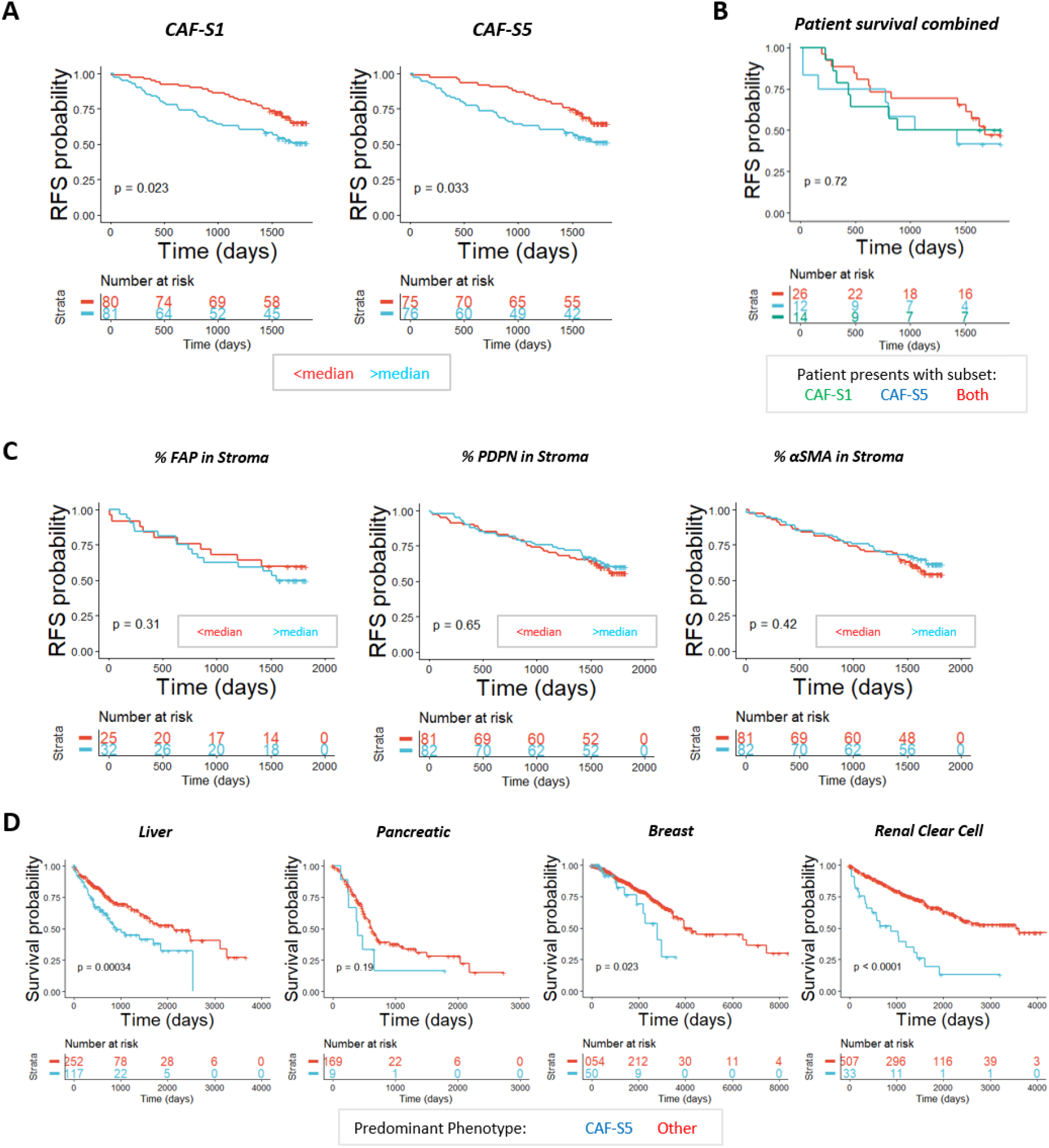
Survival analysis of CAF-S1 and CAF-S5 in NSCLC and other solid organ cancers. (A) Relapse free survival analysis of CAF-S1 and CAF-S5 when the proportion of the CAF subset present is greater or less than the median proportion expressed across all 163 patients; (B) Comparison of survival when patients only have CAF-S1 or CAF-S5 present above median levels or both, n=12 CAF-S1 only, n=26 both, n=14 CAF-S5 only; (C) Relapse free survival looking at the percentage expression of FAP, PDPN and αSMA individually in the stroma, comparing above and below median expression; (D) Survival in other cancers (hepatocellular carcinoma, pancreatic adenocarcinoma, invasive breast carcinoma and renal clear cell carcinoma) from the TCGA dataset where each patient is defined as displaying a predominant phenotype by looking at FAP (high), PDPN (high) and αSMA (low) expression.

To understand why these subsets contributed to poorer overall RFS, we investigated whether it was a single marker contributing to this by looking at the RFS of patients when the percentage of FAP, PDPN or αSMA in the stroma was above the median of all patients (Fig 6C). This revealed that there was no single marker causing such significant difference in RFS with the subsets present, although FAP did reveal a trend associated with poorer RFS with higher FAP expression.

Using the TCGA dataset we analysed the survival of patients who we expect to have greater prevalence of CAF-S5 (based on bulk high expression of FAP and PDPN in the patient and low αSMA) across four solid organ cancers: hepatocellular cancer, pancreatic adenocarcinoma, invasive breast cancer and renal clear cell cancer (Fig 6D). This revealed that the presence of these markers indicating CAF-S5 correlated with poor survival probability across these cancers.

## Discussion

Here we have identified that in NSCLC, CAFs present as a heterogeneous population which can be divided into subsets depending on their expression levels of seven fibroblast markers. Heterogeneity of CAFs is found both between and within patient samples. Two of the subsets, CAF-S2 and CAF-S3, express low levels of these markers used to identify activated fibroblasts, with CAF-S2 having low or negative expression across all markers and CAF-S3’s most significant difference from CAF-S2 being the upregulated expression of FSP1. This, and the finding that there is greater presence of these subsets in non-cancerous lung tissue compared to tumour, suggests that these subsets are representative of a more normal, healthy lung fibroblast, not one in an activated state. The other subsets identified, CAF-S1, CAF-S4 and CAF-S5 are more prevalent in tumour tissue. CAF-S5 is a novel subset, identified here as expressing FAP^Med^ CD29^Med^ αSMA^Neg-Low^ PDPN^Med^ CD90^Low^ FSP1^Low^ and PDGFRβ^Med^.

These fibroblast markers can also be used to identify CAF subsets through multiplex immunofluorescence imaging when the definitions outlined from the flow cytometry analysis are converted to binary definitions. The three subsets identified as more prevalent in the tumour (CAF-S1, CAF-S4 and CAF-S5) were investigated by staining for CAF markers FAP, αSMA, PDPN, CD90 and FSP1. Assessing the distribution of each marker across different tissue classes revealed differences between adenocarcinoma and squamous cell carcinoma, notably the expression of PDPN being higher in squamous cell carcinoma. The expression of PDPN has been linked to poor prognosis in cancer, and is hypothesised to play roles in invasion, epithelial to mesenchymal transition (EMT) and metastasis (35,36). The expression of PDPN on CAFs has been investigated in other studies, with one finding that PDPN positivity was correlated with greater invasiveness in lung adenocarcinomas (37). It would therefore be expected PDPN+ CAF subsets (CAF-S1 and CAF-S5) would be associated with poorer long-term survival, and this was indeed found in our study when assessing RFS.

Comparing the proportions of CAF subsets between NSCLC subtypes, we observed a higher proportion of CAF-S1 and CAF-S5 present in squamous cell carcinoma, and a higher proportion of CAF-S4 present in adenocarcinoma. This distribution is likely due to the expression of PDPN in CAF-S1 and CAF-S5 as previously discussed. To further characterise differences between CAF-S1 and CAF-S5, and to ensure that they were distinct populations, we analysed the single cell RNA sequencing dataset for NSCLC, published by Lambrechts et al (20). A subset of fibroblasts defined as CAF-S1 or CAF-S5 by our established criteria were compared. As the defining difference between the two subsets is the expression of αSMA, the main predicted difference was that CAF-S5 would not be a contractile phenotype. This was further confirmed by the finding that genes such as TAGLN and TPM2 were downregulated in CAF-S5, as they would contribute to contractility also, and that contractile pathways were supressed (Fig S2). The upregulation of complement genes C3 and C7 suggests that CAF-S5 are an inflammatory subset while CAF-S1 are a contractile subset.

RFS probability was found to be worse when CAF-S1 or CAF-S5 were present above median levels in NSCLC patients, despite undergoing curative resection. When considering the three markers used to identify these subsets (FAP, PDPN, αSMA), we found that in our cohort each marker did not predict RFS independently, it was only when they were considered as co-expressing in the stroma (as identified by CAF subsets) that RFS was impacted. As both CAF-S1 and CAF-S5 contribute to poor overall survival, this suggests that CAFs co-expressing FAP and PDPN are indicative of poor survival outcome in NSCLC.

To further understand the influence of the novel CAF-S5 subset on survival in other cancers we analysed the TCGA dataset for multiple solid organ cancers (liver, pancreatic, breast and renal clear cell). As this is bulk sequencing data we considered patients with increased expression of FAP and PDPN and low αSMA likely to have a dominant phenotype of CAF-S5. For these cancers there was decreased survival probability when CAF-S5 was dominant, compared to all other patients in the cohort. It has previously been shown that patients expressing the CAF-S1 phenotype in breast cancer have increased survival probability compared to groups (17). Our analysis suggests the CAF-S5 subset should be considered as a marker of poor prognosis across multiple solid organ malignancies and highlights the importance of the CAF-S5 subset as a predictor of poor outcome.

## Conclusions

We have identified five subsets of CAFs in NSCLC, including a previously undefined CAF subset CAF-S5. We have shown that CAF-S1, CAF-S4 and CAF-S5 are the most distinct to tumour tissue compared to non-cancerous tissue. CAF-S1 and CAF-S5 have been shown to be distinct populations, with CAF-S1 being FAP+, PDPN+ and αSMA+ and CAF-S5 being FAP+, PDPN+ and αSMA-, and concluding that CAF-S1 display a contractile phenotype whereas CAF-S5 display an inflammatory one. Their presence was shown to contribute to poorer overall RFS in NSCLC and suggests a poor prognosis across multiple cancer types.

## Acknowledgements

We would like to thank all the staff at the department of Thoracic Surgery, Royal infirmary of Edinburgh. We are grateful for assistance from CIR Flow Cytometry and Shared University Research Facilities, University of Edinburgh. We also thank Irene Young and Katie Hamilton for undertaking patient consents.

## Supplementary Figures

**Figure S1:**
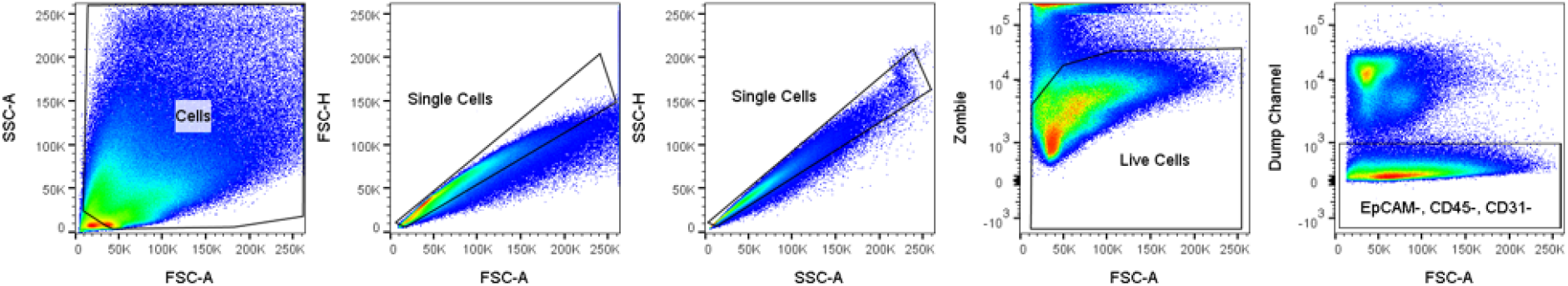
Gating strategy used to identify fibroblasts. Fibroblasts were defined as single, live cells which were EpCAM, CD45 and CD31 negative.

**Figure S2:**
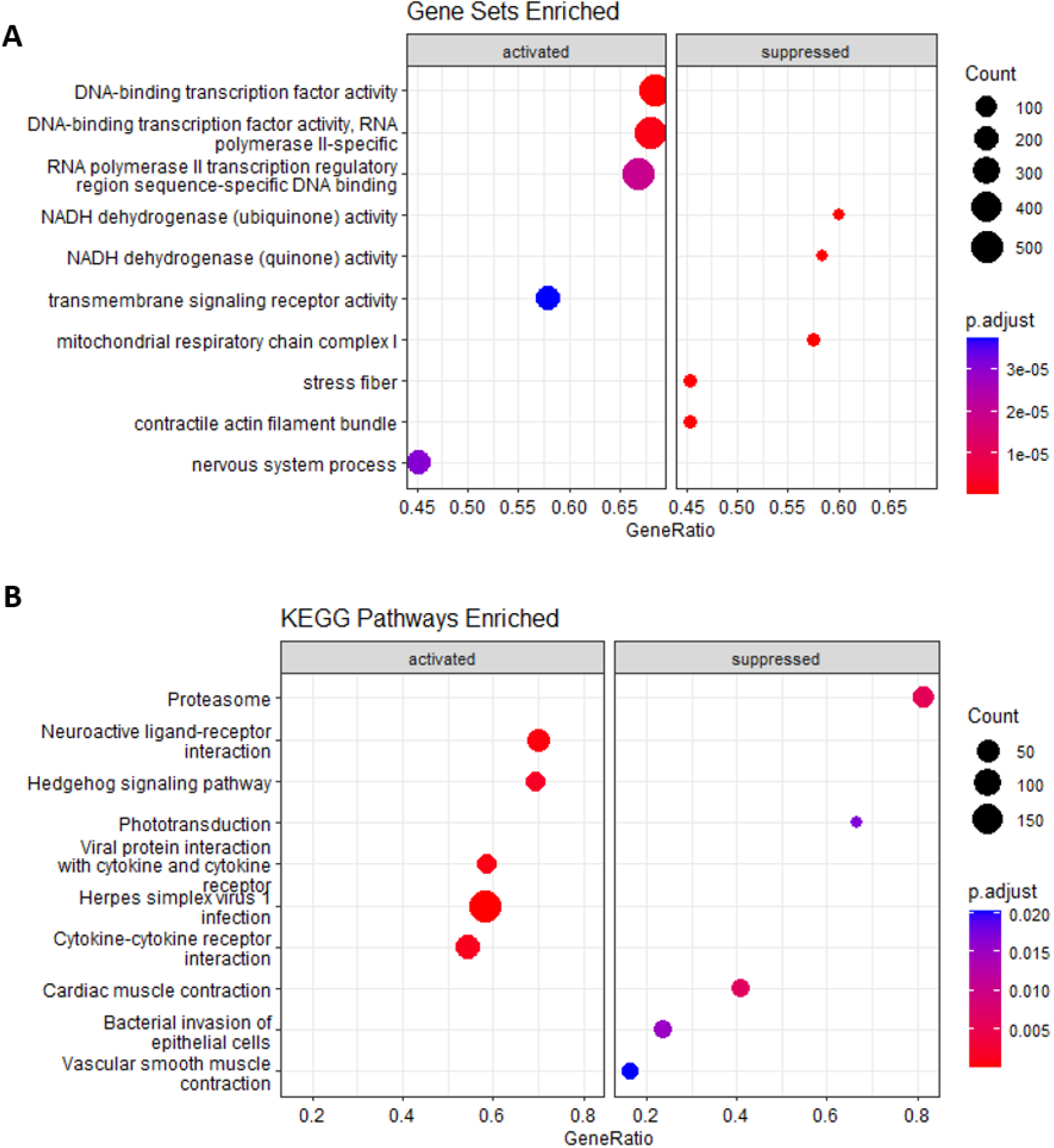
Gene set and pathways identified as enriched. (A) Gene sets activated and suppressed in CAF-S5 compare to CAF-S1 following gene set enrichment analysis; (B) KEGG pathways activated and suppressed in CAF-S5 compared to CAF-S1 following KEGG pathway analysis.

